# Ratiometric iGluSnFr imaging to assess tonic glutamate in the cerebral cortex

**DOI:** 10.64898/2026.06.12.731919

**Authors:** Moritz Armbruster

## Abstract

Tonic glutamate signaling by ambient levels of extracellular glutamate has been implicated in development, brain injury, pathologies, and physiological activity. However, it has been difficult to assay extracellular glutamate changes with spatial and temporal resolution. Here, we utilize the rarely used ratiometric excitations properties of the fluorescence glutamate sensor iGluSnFr to enable the characterization of ambient glutamate levels in acute brain slices. This ratiometric imaging enables a spatial, temporal and calibratable assay of ambient glutamate and demonstrates regional differences in ambient glutamate and sensitivity to glutamate transporters and system Xc inhibition.

## Introduction

Tonic glutamate signaling is the activation of glutamate receptors by the ambient extracellular glutamate concentration in the brain, rather than as a result of phasic synaptic glutamate release. Tonic glutamate signaling has been implicated in development (Cavelier et al., 2004; Le Meur et al., 2007; Hanson et al., 2019), following brain injury (Hinzman et al., 2010), in pathologies (Yang et al., 2019; Heit et al., 2023), and in physiological activity (Povysheva and Johnson, 2012; Balmer et al., 2021). Tonic glutamate signaling is thought to activate N-methyl-D-aspartate (NMDA) receptors especially NR2C (GRIN2C) & NR2D (GRIN2D) containing receptors (Herman and Jahr, 2007; Le Meur et al., 2007; Paoletti et al., 2013; Hanson et al., 2019) and metabotropic glutamate receptors (mGluRs)(Chen and Roper, 2004). Proposed roles for tonic glutamate signaling include modulating neuronal activity during critical periods, tonic activation, cell death, and shifting input/output relationships (Le Meur et al., 2007). Early in development, tonic glutamate levels are elevated due to lower glutamate transporter expression(Hanson et al., 2015). In adult mice, when there is robust glutamate transporter expression, tonic glutamate is mainly regulated by the System Xc-cystine/glutamate antiporter(Baker et al., 2002; De Bundel et al., 2011; Conrad and Sato, 2012).

One of the limitations in studying tonic glutamate signaling has been the challenge of assaying the extracellular glutamate concentration. Existing techniques include *in vivo* microdialysis and NMDA electrophysiological recordings. NMDA tonic glutamate assays utilize tonic NMDA currents compared to NMDA receptor inhibition and/or saturation with comparable calibration curves established with nucleated patch recordings (Herman and Jahr, 2007). The use of NMDA as an agonist bypasses the robust glutamate clearance present in the central nervous system (CNS), which restricts excitatory signaling and makes calibrations difficult(Danbolt, 2001). These recordings estimate extracellular glutamate in acute slice preparations in the low nano-molar range (∼50 nM) (Herman and Jahr, 2007; Hanson et al., 2019). *In vivo* microdialysis or glutamate biosensor voltammetry recordings of tonic glutamate involves the placement of a recording probe in the animal, which enables a slow temporal measurement of glutamate concentrations, but is limited to the spatial domain immediately surrounding the probe tip. These techniques generally report glutamate concentrations in the low micro-molar ranges (Chefer et al., 2009; Moussawi et al., 2011). Potential explanations for the order of magnitude differences in the tonic glutamate concentrations have focused on the spatial component, with NMDA recordings being biased towards synaptic sites, while microdialysis probes assay the spatial area immediately surrounding the probe tip, with potential for focal damage from the implantation. Additionally, differences are attributed to *in vitro* preparations versus *in vivo* recordings where slice preparations are continually perfused, potentially diluting their glutamate concentrations (Moussawi et al., 2011). However, both approaches are limited in their time and spatial-resolution.

In an effort to add to the toolsets available to investigate ambient glutamate, we utilized a lesser appreciated feature of the genetically encoded glutamate sensor, iGluSnFr (Marvin et al., 2013; Marvin et al., 2018) to enable ratiometric imaging of tonic glutamate in acute slice preparations of the cerebral cortex. The green fluorescent protein (GFP)-based iGluSnFr variants (iGluSnFr/SF-iGluSnFr) show an inverse response to excitation with 470nm and 405nm laser lines (Marvin et al., 2013; Marvin et al., 2018), enabling a single fluorophore excitation based ratiometric imaging of glutamate and an iGluSnFr-based assay of the resting glutamate concentration. This technique enables spatial imaging of the resting glutamate concentrations across brain regions, or sub-cellular structures, in addition to enabling a fast temporal readout in both *in vitro* and *in vivo* preparations. This complements existing techniques and sheds new light on tonic glutamate signaling.

## Materials and Methods

All animal protocols were approved by the Tufts Institutional Animal Care and Use Committee (IACUC).

### Adeno-associated virus injection

C57BL/6 male and female mice (P0) were injected intraventricullarly with high affinity SF-iGluSnFr (Marvin et al., 2018) adeno-associated virus (AAV) for the experiments in a single hemisphere. Pups were anesthetized with ice for surgery. Viruses were injected unilaterally (1μL; ventricle with ∼1 × 10^10^ gene copies per virus). Mice were housed in 12:12 light/dark cycles following surgeries and were used for acute slice preparations 40-60 days following injection. pAAV.hSynapsin.SF-iGluSnFR.A184S was a gift from Dr. Loren Looger (Addgene viral prep # 106174-AAV1).

### Preparation of acute brain slices

Cortical brain slices were prepared from AAV-infected mice as previously described (Armbruster et al., 2016). Briefly, mice were anesthetized with isoflurane, decapitated, and the brains were rapidly removed and placed in ice-cold slicing solution containing (in mM): 2.5 KCl, 1.25 NaH_2_PO_4_, 10 MgSO_4_, 0.5 CaCl_2_, 11 glucose, 234 sucrose, and 26 NaHCO3 and equilibrated with 95% O2:5% CO2. The brain was glued to a Vibratome VT1200S (Leica Microsystems, Wetzlar, Germany), and slices (400 μm thick) were cut in a coronal orientation. Slices were then placed into a recovery chamber containing aCSF comprising (in mM): 126 NaCl, 2.5 KCl, 1.25 NaH2PO4, 1 MgSO4, 2 CaCl2, 10 glucose, and 26 NaHCO3 (equilibrated with 95% O2:5% CO2). Slices were allowed to equilibrate in aCSF at 32°C for 1 hr.

### Live imaging

iGluSnFr slices were placed into a submersion chamber (Warner), held in place with small gold wires, and perfused with aCSF equilibrated with 95% O2:5% CO2 and circulated at 2 ml/min at 34°C. For stimmed experiments, a tungsten concentric bipolar stimulating electrode (FHC; Bowdoin, ME, USA) was placed in the deep cortical layers, and the upper cortical layers were imaged with a 20× water-immersion objective (LUMPLANFL, Olympus) on a custom Prior Open-Scope with X-light V2 spinning disk (Crest-optics, 89 North LDI). The 100 μs stimulus pulses were generated through a stimulus isolator ISO-Flex (A.M.P.I.). Stimulus intensity was set at 2× the resolvable threshold stimulation. Imaging was performed using a Prime95B (Photometrics), 16 bit digitization, 10 ms rolling shutter mode for 100 Hz temporal resolution, illuminated by a 470 or 405nm laser (89North) controlled by MicroManager (Edelstein et al., 2014). iGluSnFr calibration experiments were imaged in confocal mode using a 10x water immersion objective (Olympus), with differing concentrations of glutamate in Tyrodes buffer (119 NaCl, 2.5 KCl, 2 CaCl_2_, 2 MgCl_2_, 25 HEPES, 30 glucose in mM buffered to pH 7.4) were puffed using a Picospritzer II (Parker Instrumentation) with 5-10 s and ∼5 psi) using broken back patch electrodes. For wash-on experiments both infected and contralateral hemispheres were imaged using a 4X objective (Olympus) every 30s, with both channels (20ms integration 470nm laser, 50ms integration 405nm laser) with the un-infected hemisphere used to control for autofluorescence. All slices during the wash-on were aligned using the NoRMCorre algorithm (Pnevmatikakis and Giovannucci, 2017) and ratiometric iGluSnFr fluorescence quantified.

## Results

### Ratiometric iGluSnFr responses

Published characterization of the SF-iGluSnFr variants shows two distinct excitation peaks: ∼490 nm showing a large increase in fluorescence upon glutamate binding and ∼390 nm showing a smaller decrease in fluorescence upon glutamate binding (Marvin et al., 2018). The secondary peak (∼390nm) showing a inverse fluorescence response has largely been ignored, but can enable ratiometric imaging of SF-iGluSnFr. Following P0 intracerebroventricular injection of the high affinity SF-iGluSnFr (AAV1-hSyn-SF-iGluSnFR.A184S), we validated the ratiometric excitation of iGluSnFr responses. Cortical imaging of SF-iGluSnFr in response to electrically-evoked glutamate release from ascending axons in the deep cortical layers showed clear positive (470nm excitation) and negative (405nm excitation) fluorescence changes (Fig. 1A). As predicted based on the published excitation spectra, the 405nm excitation had significantly smaller magnitude ΔF/F change upon stimulation than the 470nm peak (10 stimuli 100Hz ΔF/F magnitude 0.049±0.017 vs 0.50±0.17 405nm vs 470nm, paired t-test p<0.04, N=6 slices). The peak responses of the two excitation lines were strongly correlated (R^2^ = 0.979), suggesting no changes in glutamate sensitivity, enabling its use for ratiometric imaging (Fig. 1B). The newer iGluSnFr3 sensor does not show the same negative fluorescence response (*data not shown*) likely due to the change of fluorophore from GFP to Venus. Interestingly, while the peaks of the iGluSnFr responses were highly correlated, the fluorescence decays differed significantly, with the 405nm excitation wavelength being significantly faster than the 470nm excitation wavelength (Fig. 1C-D) (Two-way Repeated Measures ANOVA, Bonferroni correction, wavelength effect p=0.011; n = 6 slices).The high affinity SF-iGluSnFr, which has a slower off-rate (Marvin et al., 2018), is not sensitive to the stimulus induced slowing of glutamate clearance as would be predicted based on the buffering of the sensor (Armbruster et al., 2020).

**Figure 1:**
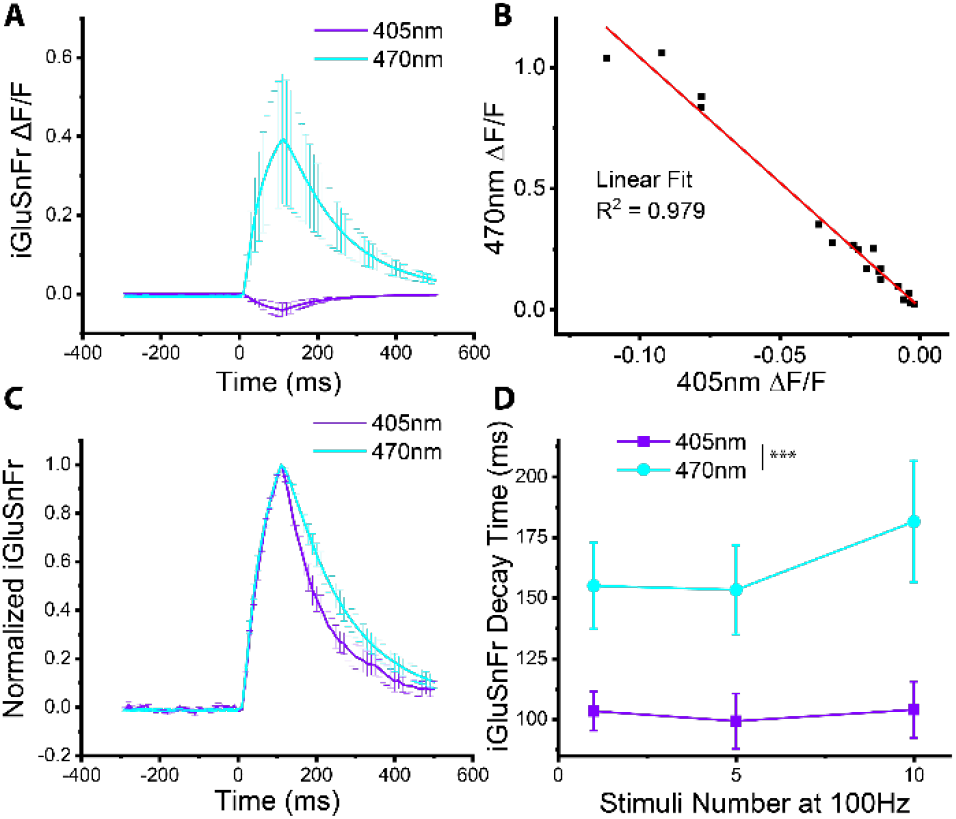
Ratiometric response of SF-iGluSnFr. A) Average fluorescence ΔF/F response of SF-iGluSnFr.A184S in the somatosensory cortex in response to 10 stimuli at 100Hz stimulation of ascending axons with 405nm or 470nm laser excitation shows opposite responses based on the excitation wavelength. B) Peak ΔF/F fluorescence (470nm; y-axis) versus (405nm; x-axis) in response to 1, 5, or 10 stimuli at 100Hz shows a strong negative correlation between the two excitation wavelengths. C) Normalized iGluSnFr responses with 405nm and 470nm excitation, suggests differences in iGluSnFr decays. D) Mono-exponential iGluSnFr decay fits to 1, 5, and 10 stimuli at 100Hz shows faster fluorescence decays for 405nm excitation (Two-way Repeated Measures ANOVA, Bonferroni correction wavelength effect p=0.011; n = 6 slices).

### Calibration

Calibrating glutamate sensors in tissue has historically been a challenge due to its excitotoxic effects and the extremely high clearance capacity of the excitatory amino acid transporter (EAAT)/glutamate transporters (Danbolt, 2001). Due to both of these reasons, it is unfeasible to simply wash on different glutamate concentrations. In order to calibrate the ambient glutamate concentration, we locally puffed different glutamate concentrations (0.1, 1, 10, 100, 1000, 10000nM) and calculated the fluorescence changes, similar to a no-net flux method used in microdialysis (Moussawi et al., 2011). We assumed these puffs will not fully clamp all the imaged iGluSnFr sensors to the puffed concentration, but instead will provide a relative change from the resting glutamate concentration to the puffed concentration. The resulting responses were then fit with a Hill equation with the cooperativity (n = 1) and affinity of the sensor (600nM) (Min = 0.96 F_glu_/F_0_, Max = 1.32 F_glu_/F_0_, R^2^ 0.98, n = 6 mice). This results in a resting glutamate concentration of 65nM with a 90% confidence interval of 2-128nM (Fig. 2). A limitation of this calibration approach is that it cannot be mapped onto other measures of the ratiometric sensor due to the no-net flux method, apparent from the reduced F_glu_/F_0_ range.

**Figure 2:**
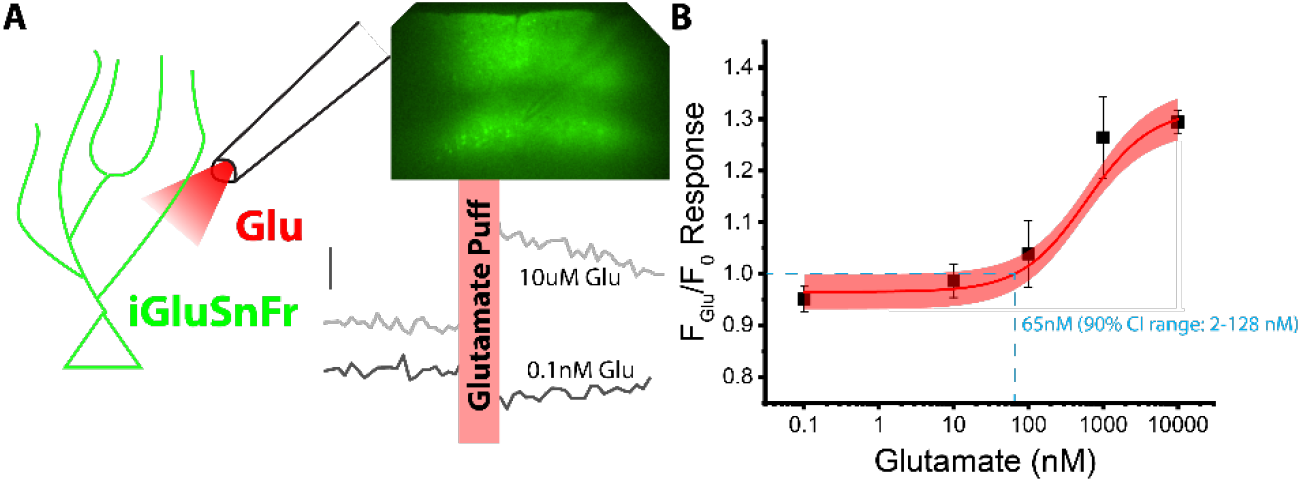
Calibration of tonic glutamate levels. A) Focal puffing of differing glutamate concentrations from a puffer pipette in the somatosensory cortex evokes increases (10 μM glutamate example trace) or decreases (0.1 nM glutamate example trace) of 470 nm/405 nm iGluSnFr ratio ΔF/F. B) Average ratiometric iGluSnFr response to fixed glutamate concentrations fit with a sigmoidal Hill equation shows resting cortical glutamate concentration at 65 nM (90 CI range 2-128 nM). n = 26 slices from 6 mice.

### What drives tonic glutamate?

This assay of ambient glutamate enables us to apply pharmacology to test the contributions of specific sources to the level of ambient glutamate. Low magnification ratiometric iGluSnFr imaging was performed of the infected hemisphere and the uninfected contralateral hemisphere as control. For all images, auto-fluorescence was subtracted from contralateral hemisphere before calculating the iGluSnFr ratio. Following a baseline, TTX (1μM) was washed in to block neuronal activity (Fig. 3), which had no effect on the ambient glutamate level(iGluSnFr ratio 2.07 vs 2.12 control vs. TTX N = 16 slices/7 mice p = 0.14 paired t-test). Subsequently, the glutamate transporter GLT-1 inhibitor DHK (75μM) or the system xc transporter sulfazlazine (100μM) was applied. Lastly, the pan glutamate transporter inhibitor TFB-TBOA (1uM). This showed that inhibiting GLT-1 led to a significant, but modest increase in the ambient glutamate concentration (Fig. 3)(iGluSnFr ratio 2.32 vs 2.54 for before vs. post DHK, N = 7 Slices/3 mice p = 2.13E-4 paired t-test). Inhibiting system xc, showed a significant increase in the iGluSnFr signal(iGluSnFr ratio 3.31 vs. 5.52 for Control vs. Sulfasalazine, N = 13 slices/7 mice p = 0.011 paired t-test). TFB-TBOA caused the iGluSnFr signal to saturate and led to a large increase (Fig. 3)(iGluSnFr ratio 2.44 vs. 7.84 Control vs TFB-TBOA N = 26 slices/7 mice p = 3.4E-5 paired t-test).

**Figure 3:**
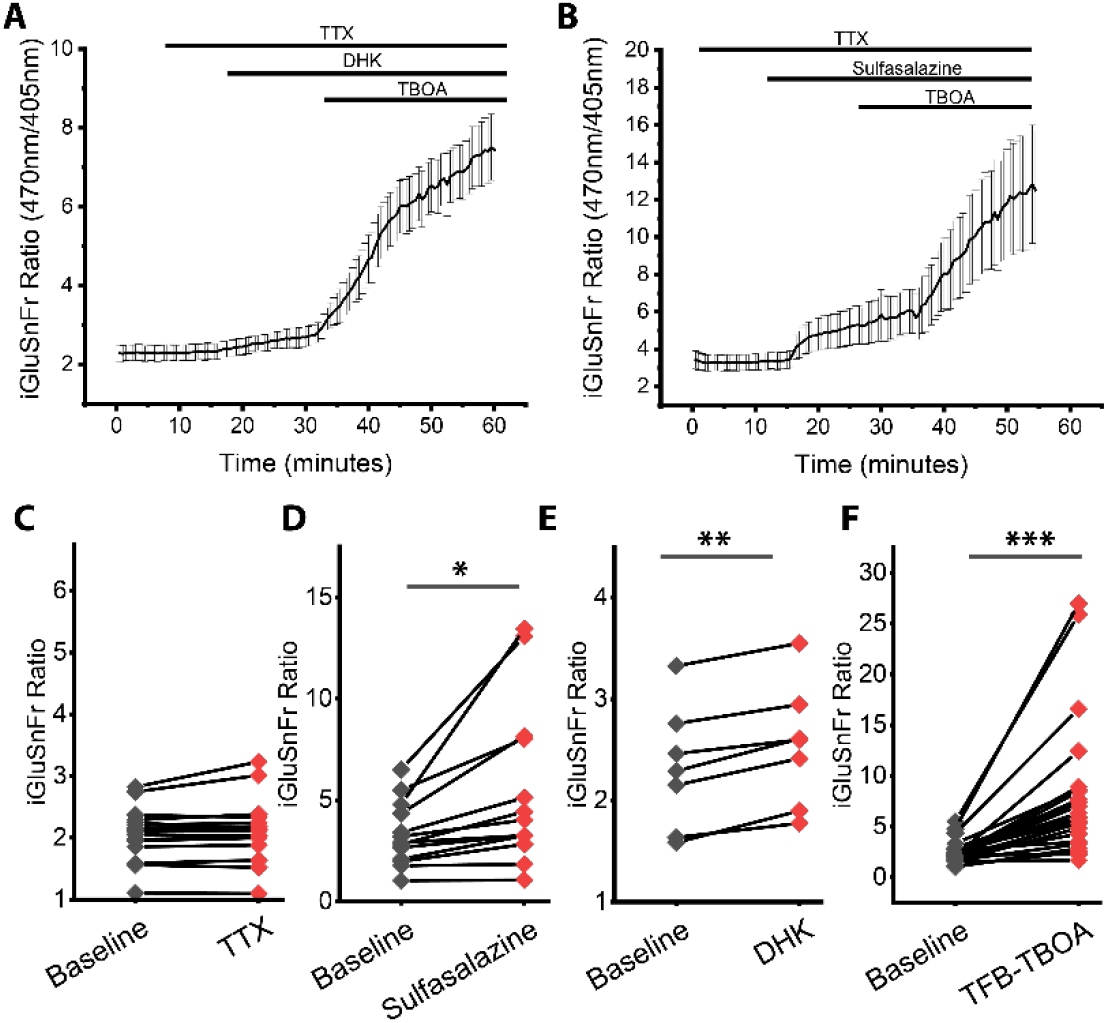
Glutamate transporters and System Xc contribute to tonic glutamate. A) Average ratiometric iGluSnFr imaging from the somatosensory cortex with TTX (1μM), DHK (75μM), and then TFB-TBOA (1μM) shows increase in iGluSnFr ratio suggesting increased ambient glutamate. B) Average ratiometric iGluSnFr in response to TTX, Sulfasalazine (100μM), and TFB-TBOA shows increases in iGluSnFr ratio in response to System Xc inhibition and the pan-glutamate transporter inhibitor TFB-TBOA. C). TTX shows no effect on iGluSnFr ratio. D) System Xc inhibitor Sulfasalazine, E) GLT-1 inhibitor DHK and F) and pan glutamate inhibitor TFB-TBOA show increases in iGluSnFr ratio (Paired t-tests; * = p < 0.05, ** = p < 0.01, *** = p < 0.001. N=16 (TTX), 13 (Sulfasalazine) 7 (DHK), 27 (TFB-TBOA) slices from 4-7 mice).

### Spatial Heterogeneity

This assay enables the assaying of ambient glutamate across different spatial domains. Here we quantified the resting iGluSnFr ratio in the cortex, hippocampus, and striatum (Fig. 4). The striatum showed a significantly higher 470nm/405nm iGluSnFr ratio compared to the cortex (iGluSnFr ratio 2.27, 2.68, 3.33 for Cortex, Hippocampus, and Striatum with N = 11, 6, 10 slices N = 4, 3, 3 mice respectively. p = 3.1E-4 Cortex vs Striatum One way ANOVA with Tukey post-test), suggesting an increase in tonic glutamate in the striatum. To further confirm this finding, the glutamate transporter inhibitor TFB-TBOA (as in Fig. 3) was washed in. If tonic glutamate is increased in the striatum, TFB-TBOA should have a smaller effect than in the cortex. Consistent with this, the striatum showed a reduced TFB-TBOA/Baseline ratio compared to the cortex, consistent with a higher resting ambient glutamate level (TFB-TBOA/Baseline ratio 3.31 vs 2.79 for Cortex vs. Striatum N = 9 slices from 3 mice, p = 0.036 paired t-test).

**Figure 4:**
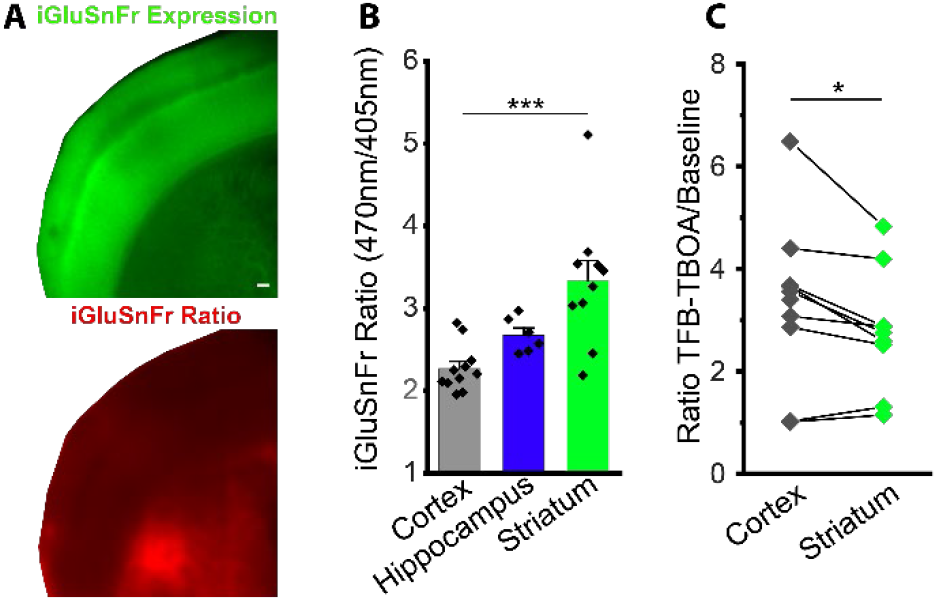
Spatial heterogeneity in tonic glutamate. A) Example image iGluSnFr expression and ratio (470nm/405nm) of cortex and striatum suggests increased resting glutamate in the striatum. B) Ratiometric iGluSnFr imaging baseline from Cortex, Hippocampus, and Striatum shows significantly higher iGluSnFr signal in the striatum (One-Way ANOVA, Tukey post-hoc test p<0.0004, N= 11, 6, 9 slices from 4, 3, 3 mice). C) Striatum iGluSnFr signal shows a smaller response to TFB-TBOA wash-on compared to the cortex, in agreement with increased tonic glutamate (Paired t-test p<0.04; N= 9 slices from 3 mice).

## Discussion

We showed that high affinity iGluSnFr variants can be utilized in a ratiometric imaging fashion to measure ambient glutamate levels. Our measured ambient glutamate level is similar to values achieved with NMDA wash-on normalizations (∼50nM) (Cavelier et al., 2004; Hanson et al., 2019) (Herman and Jahr, 2007) (Le Meur et al., 2007) and on the very low end of values reported with microdialysis (Turecek and Trussell, 2000)(Benveniste et al., 1984; Masse et al., 2019). This is in contrast to higher ambient glutamate levels often reported by microdialysis (Chefer et al., 2009; Moussawi et al., 2011) or glutamate voltammetry probes. Differences between these assay methods have been previously discussed including *in vitro* versus *in vivo* preparations, potential localized glutamate domains, and damage due to probes or slice preparations. This ratiometric iGluSnFr assay adds to the existing toolset to measure ambient glutamate with the additional benefits of enabling spatial and temporal resolution Our iGluSnFr based method of recording the ambient glutamate levels has a number of potential advantages compared to existing approaches. These include combining both the temporal and spatial resolution of an imaging approach and the ease of measurement compared to NMDA normalizations. This is highlighted by our current results suggesting higher ambient glutamate levels in the striatum compared to the cortex. This finding is consistent with a previous microdialysis approach showing increased tonic glutamate in the striatum compared to the prefrontal cortex (Baker et al., 2003) and is supported by the striatum having slower glutamate clearance kinetics than the cortex or hippocampus (Pinky et al., 2018). However, with some exceptions (Moghaddam, 1993; Bagley and Moghaddam, 1997), there has been few comparisons of ambient glutamate levels across or within brain regions, something that is more easily assayed with a spatial fluorescence indicator. Also of note is the variability in the baseline iGluSnFr ratio, with a large spread of the 470nm to 405nm fluorescence ratio across slices (Fig. 3). It is unclear if this range is due to the sensor (trafficking, expression levels), differences in glutamate, and/or other factors and raises a concern about comparing baseline levels across slices or between slices unless secondary correlative measures such as TFB-TBOA wash-on can also be utilized.

We expect that this imaging approach should be compatible with *in vivo* preparations as well. However, there are two limitations of our current study: the affinity of the already high-affinity iGluSnFr sensor is at the limits of its sensitivity for the resting glutamate concentrations observed and calibrating the sensor is difficult as we can not clamp the glutamate concentration. Ideally, an even higher-affinity sensor that is compatible with fluorescence lifetime imaging would be ideal. Promisingly, some iGluSnFr3 variants show potential for fluorescence lifetime imaging (Aggarwal et al., 2023). This approach could also be utilized with appropriate high-affinity sensors, to other signaling molecules such as GABA (iGABASnFr2), dopamine (dLight)(Patriarchi et al., 2018), neuromodulators D-serine (Vongsouthi et al., 2021), and acetylcholine (Jing et al., 2018), among others. Which would open new possibilities, including investigating spatial heterogeneity of these signaling molecules within and across brain regions.

Microdialysis approaches suggests that ongoing activity (TTX-sensitive) does not significantly contribute to ambient glutamate levels; however, glutamate biosensor voltammetry does show a significant TTX-sensitive component. Our iGluSnFr imaging data mirrors the microdialysis results and does not show a significant TTX sensitive component, suggesting that on-going neuronal activity does not significantly contribute to ambient glutamate concentrations. This is in-line with studies that suggest glutamate is captured within milliseconds of release by glutamate transporters surrounding synapses (Armbruster et al., 2020). Microdialysis studies suggests that System Xc contributes significantly to setting the ambient glutamate concentration by exporting glutamate with the import of cystine (Lewerenz et al., 2013). Our data surprisingly shows an increasing iGluSnFr signal with the inhibition of System Xc, suggesting an increase in ambient glutamate. A number of differences exist in our preparation compared to the microdialysis finding of glutamate export via System Xc, including, *in vitro* acute slices versus *in vivo* recordings, differences in ongoing activity levels, and differences in basal ambient glutamate levels. However, little evidence in the literature supports reversal of the System Xc transporter, and as such, we suspect that our increase in glutamate levels following System Xc inhibition may be due to potential secondary effects such as antioxidant-induced swelling inhibiting EAATs (Jayakumar et al., 2009; Li et al., 2022). In conclusion, we have shown that the GFP based iGluSnFr reporters can be used via ratiometric excitation to quantify ambient glutamate levels spatial across regions and temporally across pharmacological application.

## Acknowledgments

This work was funded by the grants from the National Institute of Neurological Disorders and Stroke (R01NS127819 to M.A.). We would like to thank Dr. Loren Looger and the GENIE Project for making plasmids and viruses available that were used in this study. Lastly, we would like to thank members of the lab for critical reading of the manuscript.

